# ROCK Inhibitor Increases Proacinar Cells in Adult Salivary Gland Organoids

**DOI:** 10.1101/712877

**Authors:** Matthew Koslow, Kevin J. O’Keefe, Deirdre A. Nelson, Melinda Larsen

**Affiliations:** Graduate program in Molecular, Cellular, Developmental, and Neural Biology, University at Albany, State University of New York, Albany, NY 12222 USA; Department of Biological Sciences, University at Albany, State University of New York, Albany, NY 12222 USA; RNA Institute, University at Albany, State University of New York, Albany, NY 12222 USA

**Keywords:** Salivary Gland, Salisphere, Organoid, Proacinar Differentiation, ROCK

## Abstract

Salisphere-derived adult epithelial cells enriched for progenitor cells have been used to improve saliva production of irradiated mouse salivary glands. Importantly, optimization of the cellular composition of salispheres could improve their regenerative capabilities. The Rho Kinase (ROCK)^1^ inhibitor, Y27632, has been used to increase the proliferation and reduce apoptosis of progenitor cells grown in vitro. In this study, we investigated whether Y27632 in different cell media contexts could be used to improve expansion of adult epithelial progenitor cells or to affect their differentiation potential. Application of Y27632 in medium used previously to grow salispheres promoted expansion of Kit^+^ cells, while in simple serum-containing medium Y27632 increased the number of cells that expressed the K5 basal progenitor marker. When tested in a 3D organoid assay, Y27632 enhanced the contribution of adult salispheres to salivary organoids expressing the secretory proacinar marker Aquaporin 5 (AQP5) in response to FGF2 dependent mesenchymal signals. Optimization of epithelial-mesenchymal interactions organoids can be used to improve application of adult salivary progenitor cells in regenerative medicine strategies.

**Highlights:**

- Y27632 promotes Kit^+^ salisphere cell proliferation in salisphere media.
- Y27632 promotes K5 expression in salispheres cultured in serum containing media.
- Y27632 treated Kit^+^ salispheres form salivary organoids expressing AQP5.

## 1. Introduction

The loss of salivary gland function, or salivary hypofunction, is a clinical condition that occurs in patients diagnosed with Sjögren’s Syndrome and in patients that have undergone radiation therapy for head and neck cancer. Currently there are 500,000 patients who suffer from this condition (Lombaert, 2017). Salivary hypofunction results in the sensation of “dry mouth” or xerostomia and leads to a decline in quality of life as saliva is essential for digestion, mineralization of teeth, lubrication of the oral cavity and immunity against microorganisms, fungi, and viruses (REFS). There are commercially available supplements to increase moisture in the oral cavity including synthetic saliva, stimulants such as pilocarpine, and moisturizers. Despite the availability of these substances, they only provide temporary relief and are not a cure. To provide permanent salivary gland functional restoration after xerostomia, progenitor cell therapies hold great promise (Lombaert et al, 2008; Nanduri et al, 2014). To develop future cell-based salivary gland regenerative therapies, increased knowledge of how to expand progenitor cells and defining the inputs that regulate their growth and differentiation are needed.

Stem and progenitor cells elaborate organs during development and are of interest for application in regenerative therapies. In the salivary gland, multiple epithelial progenitor cell populations have been identified, and the relative contribution of each cell type during development and during injury repair remains a topic of great interest. During the later stages of branching morphogenesis cells in the end buds that will become secretory acinar cells start to differentiate. mRNA for the water channel protein Aquaporin 5 (AQP5) begins to be expressed in the end buds by embryonic day 14 (E14), with AQP5 protein expression detectable the following day E15 with membranous localization occurring by E16 (Nelson et al, 2013). In adults, acinar cells replicate through a process of self-duplication rather than through differentiation from stem or progenitor cells (Aure et al, 2015). Multiple progenitor cell populations that replenish the ducts reside in the SMG, including cells which express the intermediate filaments cytokeratin 5 (K5) and cytokeratin 14 (K14), which are basal progenitor markers in other branching organs, including the mammary gland (Rios et al, 2014). K5^+^ cells can differentiate into more mature lumenal ductal cells that express Prominin 1 (DeSantis et al, 2017). K5 differentiation is controlled by parasympathetic innervation (Knox et al, 2010) and through retinoic acid receptor (RAR) signaling (Abashev et al, 2016; DeSantis et al, 2017). Kit is a receptor tyrosine kinase that marks progenitor cells in many different cell types but is reported not to be a stem cell marker in adult salivary glands (Kwak et al, 2018; Emmerson et al, 2018). Kit^+^ cells are found in end buds in developing glands, but Kit^+^ cells transition primarily to the ducts after birth (Nelson et al, 2013; Wang et al, 2014). Kit^+^ cells isolated from salispheres partially restored saliva production after implantation into irradiated SMGs (Lombaert et al, 2008; Pringle et al, 2016). While progenitor characterization has been well documented in the context of embryonic SMG development, there is a lack of understanding on which SMG progenitors contribute to proacinar differentiation in organoids and how they function in cell therapy for xerostomia.

A potential method to enhance progenitor cell expansion in salisphere cultures is through manipulation of the Rho-associated protein kinase (ROCK) pathway. ROCK is a member of the serine-threonine kinase family that is activated by the small GTPase, RhoA (Amamo et al, 2010), and has many functions in the cell, including promoting apoptosis (Coleman et al, 2001). Due to these properties, ROCK inhibition has been used to stimulate proliferation and prevent apoptosis of stem/progenitor cells during expansion in culture. ROCK inhibition by Y27632 has been shown to increase survival of dissociated human embryonic stem cells and to promote expansion of tissue resident stem cells in culture (Watanabe et al, 2007. Zhang et al, 2011). ROCK inhibition has been shown in to increase progenitor expansion in many cell types (Kim et al, 2015; Zhou et al, 2016). ROCK signaling is critical for salivary gland development (Daley et al, 2009, 2011, 2012; Gervais et al 2016), and ROCK inhibition has been shown to delay senescence and promote proliferation of the epithelial cells from adult SMG (Han et al, 2018; Nanduri et al, 2014; Lee et al, 2015). However, a role for ROCK inhibition to promote expansion of specific progenitor cell populations in adult derived salispheres has not been investigated.

Organoids are self-assembled 3D collections of cell types that mimic the morphology and functionality of full organs at a smaller scale *in-vitro*. Single progenitor cells or tissue pieces can be used to generate organoids (Kretzschmar and Clevers, 2016). When specific factors are added *in vitro*, organoids are able to self-organize into structures that resemble mini organs (Rossi et al, 2018; Lancater and Knoblich, 2014). Significantly, implantation of organoids into damaged or diseased organs has been shown to improve functionality of several adult organs in animal models, and organoids are important models to study cell interactions that control differentiation and tissue organization in vitro (Karthaus et al, 2014; Kretzschmar and Clevers, 2016; Ramachandran et al, 2015; Tan et al, 2017). We previously established methods to form salivary gland organoids from primary embryonic cells (Hosseini, 2018a,b). We used organoids to identify a requirement for FGF-2 signaling in the mesenchyme and laminin-111 for development of complex, branched proacinar organoids from embryonic E16 epithelium (Hosseini et al, 2018 a,b). In this study, we investigate the contribution of ROCK signaling in organoid formation. We show that the ROCK inhibitor, Y27632, and different media compositions can be used for improved expansion of defined mouse adult SMG progenitor cell populations in salisphere cultures and that salispheres expanded with Y27632 have an increased capability of generating complex FGF2-induced proacinar salivary organoids.

## 2. Materials and Methods

### 2.1. Mouse Strains

CD-1 (Charles River) or C57BL/6 (JAX #000664), adult (8-12 weeks old) or timed-pregnant female mice were obtained from Charles River Laboratories, and are used as indicated in the figure legend. *Mist1*^*CreERT2*^ mice were gifted to us by Dr. Catherine Ovitt from University of Rochester with permission from Dr. Steven Konieczny, Purdue University. *Mist1*^*CreERT2*^ mice were bred to the ROSA26^*TdTomato*^ reporter strain (JAX #007909), and genotyped by PCR to detect Cre and TdTomato, as previously described (Le and Saur, 2000; Inoue et al, 2010; Fujiwara et al, 2016). Mice were induced with tamoxifen (Sigma cat# T5648) (1 mg/ml) dissolved in corn oil and ethanol, injected intraperitoneal (IP) with one 5mg/kg of body weight dose delivered every other day for three days, and tissue harvested one week from the first induction. 5 mg of tamoxifen was dissolved in 450 µL of corn oil and 50 µL of 100% v/v ethanol (10 mg/ml stock) to make 1 mg/ml stock solution. To confirm success of tamoxifen injection, RFP fluorescence of live SMG tissue was examined with an inverted fluorescent microscope (EVOS, Invitrogen). All CD-1 or C57BL/6mice were aged 7-11 weeks prior to harvest. *Mist1*^*CreERT2*^; ROSA26^*TdTomato*^ mice were aged 18 months. SMG from female mice were primarily used but males were used as indicated in specific experiments. All animals were maintained and handled using protocols approved by the University at Albany Institutional Animal Care and Use Committee (IACUC).

### 2.2. Salivary Gland Removal and Primary Cell Isolation

E16 mouse SMGs were dissected by removing a mandible slice with a scalpel and removing the glands from the slice with sterile forceps under a dissecting microscope. Once E16 salivary glands are harvested, they are processed as previously described (Hosseini et al, 2018 a,b). For adult glands, cells were similarly digested as with embryonic tissue with the addition of 30 µg/mL DNAse I for 10 minutes. (Stem Cell Technologies #07900). Time zero (T0) gland preparations were prepared as follows. After digestion, cells were mixed in a 1:1 mixture of cells in Media to Matrigel®. 10 µL of this mix in placed on a Nuclepore filter. Cells are seeded at 10^5^ per filter and incubated for 30 minutes at 37C. These cells are immediately fixed and prepped for ICC as described below. In addition, epithelium went through an adhesion depletion stage in a 35 mm dish for two hours to reduce contaminating stromal derived cells as previously reported (Hosseni et al, 2018a).

### 2.3. Preparation of enriched primary E16 SMG mesenchyme for Organoid Assay

Primary mesenchyme cells were isolated as previously described Hosseini 2018b). Briefly, E16 SMG were treated with enzymes to release epithelial clusters and subjected to gravity sedimentation. Cells in the supernatant were filtered through a 70 µM filter (Falcon REF 352350) followed by a 40 µM filter (Fisherbrand 22363547). To deplete EpCAM^+^ epithelial cells, EpCAM microbeads (130-105-958, Miltenyi Biotech) were incubated with the cell preparation and collected with the MACS sorting column, as previously described (Kwon et al, 2017).

### 2.4. Culture Media

Simple serum-containing medium was made from Dulbecco’s Modified Eagle Medium: Nutrient Mixture F-12 (DMEM/F12) (Life Technologies 11039047) supplemented with 100 U/mL penicillin, 100 mg/mL streptomycin (Pen-Strep, Life Technologies), and 10% v/v Fetal Bovine Serum (FBS) (Life Technologies # 10082-147). “Salisphere” (Sali) medium is made with DMEM/F12 together with the following components: 100 ng/mL Fetal Growth Factor-2 (FGF2 Peprotech #450-33), 100 ng/mL Epidermal Growth factor (EGF, PeproTech #AF100-15), 10 µg/mL Insulin (Sigma #I882), 1.25 µg/mL Hydrocortisone (Sigma #H0135), 1% v/v N2 Serum Supplement, L-Glutamate, and Pen-Strep (Life Technologies # 15140122). Y27632 was obtained from Sigma-Aldrich (CAS 146986-50-7) dissolved in DMEMF12 media and stored as frozen single-use aliquots. Y27632 was added to the culture media for a concentration of 10 µM.

### 2.5. Salisphere Culture

Epithelial clusters derived from two adult glands were seeded in 2 mL of media in a 6-well dish at 2×10^6^ cells per well (approximately 0.5 glands/well). Adult epithelium was cultured for three days to generate salispheres. At 72 hours, the salispheres were sedimented by gravity and subjected to adhesion depletion to remove stromal-derived cells. 1000 salispheres were seeded on top of 0.1 μm pore porous polycarbonate filters (Nuclepore, Whatman #0930051) and floated on top of 200 µL media (control or control media with 100ng/ml FGF2) in 50 mm glass-bottom dishes (MatTek P50G-1.5-14F) for 7 days.

### 2.6. Organoid Assay

Salispheres were collected by media removal, and 10 µl of cold of 1:1 media: Matrigel® added to the clusters and gently pipeted. Where indicated, 2*10^4^ primary mesenchyme cells derived from E16 SMGs were mixed with 1000 salispheres or to 1.25 glands worth of E16 epithelium (2.75*10^5^ cells) in either control media or 100 ng/mL FGF2. Salispheres were cultured for 7 days unless otherwise specified and media was changed every 72 hours to form organoids.

### 2.7. ICC and Confocal Imaging

Salispheres are either fixed in 4% v/v Paraformaldehyde (PFA) (Electron Microscopy Sciences) at 4 degrees overnight or 100% v/v Methanol (simultaneous fix/permeabilize) at −20 degrees for 18 minutes. PFA-fixed cells for ICC were performed as described previously (Hosseni et al, 2018 a,b), secondary antibody incubations were performed at room temperature for 2 hours. Cells were then blocked in a mix of 3% v/v BSA in PBS with 5% v/v donkey serum and Mouse on Mouse (M.O.M) blocking agent (Vector Laboratories MKB-2213), for 2 hours. The primary antibodies were utilized as follows. FITC-EpCAM (1:200, eBiosciences #11-5791-82), Vimentin (1:1000, Sigma #V2258), cytokeratin 5 (K5) (1:200 BioLegend # 905501), cytokeratin 14 (K14) (1:400 BioLegend # 906001), AQP5 (1:200. Alomone #AQP-005), c-Kit (1:200 R&D Systems #AF1356) Ki67 (1:200 AbCAM), CC3 (1:200, #Cell Signaling 9661), RFP (1:200, Abcam #ab62341) with DAPI (Life technologies # D1306 1:1000) to label nuclei. Secondary antibodies including Cyanine and Alexa dye-conjugated AffiniPure F(ab^’^)_2_ fragments were all purchased from Jackson ImmunoResearch Laboratories and used at a dilution of 1:200. Nuclepore filters were mounted on glass slides using Fluorogel with a Tris buffer mounting media (Electron Microscopy Sciences #17985-11). Images for quantification were acquired using the Zeiss Cell Observer inverted fluorescent microscope. Confocal microscopy was performed using a Zeiss LSM710 laser scanning confocal microscope at 20X magnification maximum projections.

### 2.8. Image Analysis and Statistical Analysis

Image J was used to perform quantification of widefield fluorescent images as previously reported (Gervais et al, 2015; Hosseni et al, 2018). The percent of cells positive for a specific marker was determined by counting the # of cells positive for the marker relative to number of cells expressing DAPI. To quantify the intensity of a marker expressed by cells, images were normalized to levels of EpCAM. Area was quantified similarly but normalized to DAPI. For statistical analysis one way ANOVA was carried out for multiple comparisons were carried out using Vassar-Stats with Tukey post-hoc test with p* < 0.05 considered to be minimally statistically significant for comparisons within the ANOVA. Student’s two-tailed T tests for 2 way comparisons were carried out as indicated. At least three experiments were quantified to generate each graph with 3 samples, unless specified. Graphs of quantified data were made using Excel. Error bars are standard error of the mean (SEM). ImageJ was also used to generate maximum intensity projections of confocal images.

## 3. Results

### 3.1. Effect of Y27632 treatment on salisphere formation and proliferation

We compared the survival and growth of epithelial cell clusters isolated from adult submandibular salivary glands as salispheres in a simple serum-containing media (DMEM:F12 +10% (v/v) FBS) that we used previously to culture embryonic epithelial clusters as organoids (Hosseini 2018 a,b) and a complex serum-free media that was previously used to grow adult mouse salispheres (Sali media) (Nanduri, et al 2011) both with and without the ROCK selective inhibitor Y27632. The number of salispheres present after three days of culture in simple media was 1503±324 while the number was nearly double when Sali media was used with 3475±345 (**Fig 1a-b**). With the addition of Y27632 there was a small increase in the number of salispheres (1783±422 in Simp+Y27632 vs 4052±159 in Sali+Y27632). We also measured the average size of the salispheres grown under different media conditions. Although there was little difference in the size of salispheres with Y27 treatment in simple medium, salispheres were larger in Sali media relative to simple media and slightly larger in Sali media +Y27 than in Sali media alone (**Fig 1 c-d**). We then examined the effect of Y27632 on the proliferation and apoptosis of salispheres. We performed ICC to detect Ki67, a protein that is expressed during all phases of the cell cycle except for G_0_, in salisphere cultures (Scholzen and Johannes, 2000; Nguyen et al, 2018). Comparing salispheres grown in simple media with salispheres grown in Sali media for three days, the number of cells expressing Ki67 was twice as high in Sali media. The inclusion of Y27632 increased the number of Ki67 positive cells grown in both medias with Y27632 having a greater impact in Sali media than in simple media (**Fig 1e-f**). Next, we examined the levels of apoptosis in cultured salispheres. ICC for the apoptosis marker, Cleaved Caspase 3 (CC3) revealed that salispheres cultured in simple medium were highly apoptotic, while treatment with Y27632 significantly reduced CC3 levels after 3 days (0.3±0.08 vs 1±0.2) (**Fig 1g-h**). In the Sali media alone, apoptosis levels were very low and Y27632 did not significantly affect apoptosis levels in this medium. Together, these results indicate that Y27632 treatment primarily promoted cell cycle entry of cells in salispheres cultured in Sali media but promoted both cell cycle entry and reduction of apoptosis in salispheres cultured in simple media.

**Figure 1:**
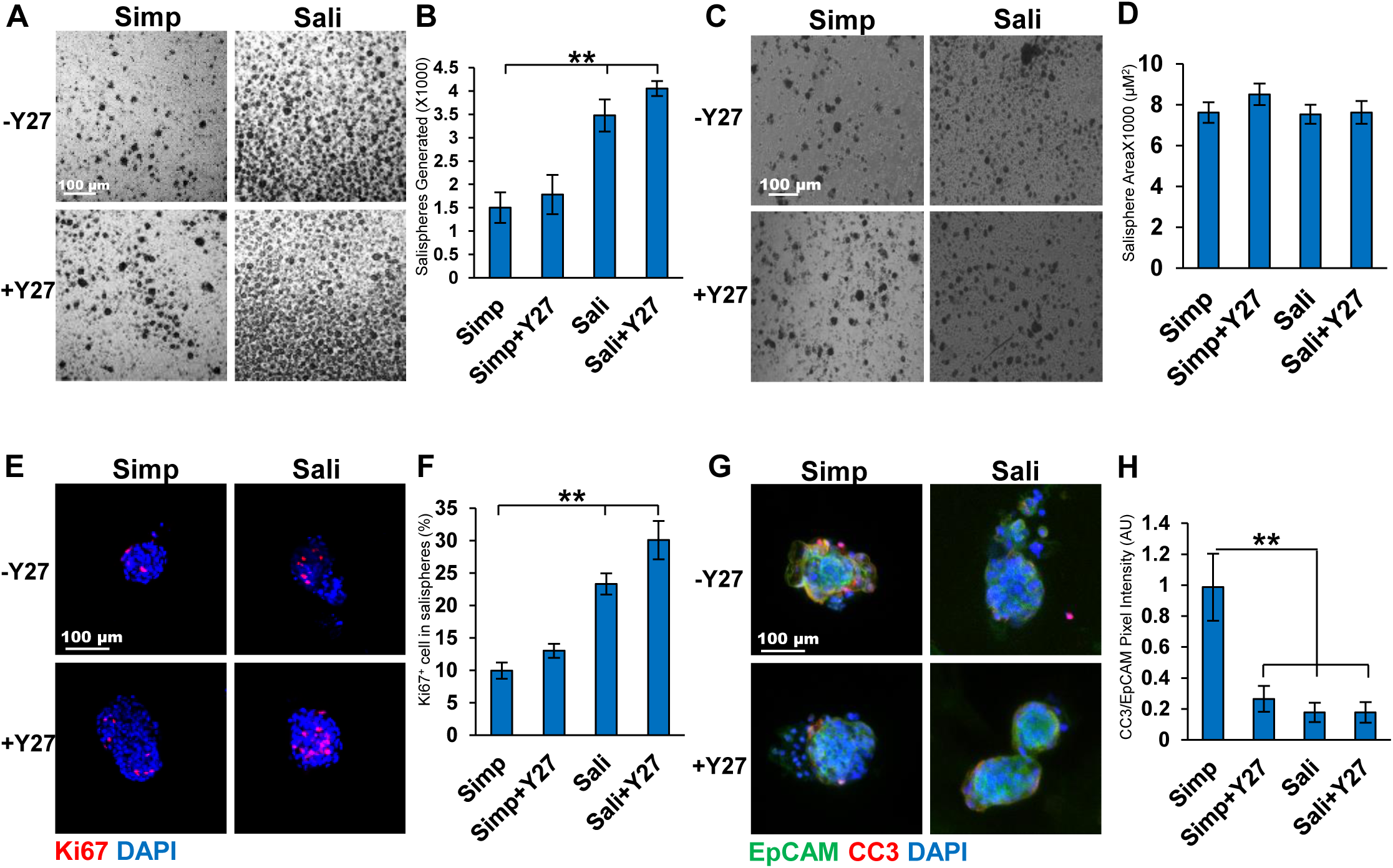
Y27632 treatment increases cell proliferation and reduces apoptosis within salispheres. A) Brightfield images of salispheres cultured for three days in Simple (Simp) or salisphere (Sali) medium +/- Y27632 (Y27). B) Number of salispheres generated at 3 days. C) Bright field of salispheres generated per gland per media condition at 3 days D) Average size of salispheres quantified at 3 days. N=6. E) ICC of Ki67 in salispheres cultured for 3 days with Ki67 (red, proliferating cells) and DAPI (blue, nuclei). F) Percent of Ki67^+^ cells per salisphere to total cells. G) ICC of Cleaved Caspase 3 (CC3, red) vs EpCAM (green, epithelium) and DAPI (blue). H) Quantification of CC3 pixel intensity relative to EpCAM. A-D: N=6, E-F: N=3. ** p < 0.01, One way ANOVA.

### 3.2. Cellular composition of salispheres ± Y27632

As Kit^+^ cells are present salispheres and of therapeutic interest in salivary gland regeneration (Lombaert et al, 2008; Nanduri et al, 2014; Pringle et al, 2016) we examined the level of Kit expression in our salispheres. Salispheres were cultured for 3 days and subjected to ICC to detect Kit and the epithelial cell surface marker, EpCAM. Kit was significantly increased in Sali Media with Y27632 relative to Sali media alone when the Kit^+^ area of the salisphere was quantified (11.6±5 vs 42.4±11), and when the intensity of Kit staining normalized to EPCAM levels was quantified (0.123±0.03 vs 0.399±0.08) (**Fig 2a-c**). Indeed, compared to samples fixed after preparation of epithelial clusters at the beginning of salisphere culture, Kit levels/intensity were fully restored in Sali+Y27632 treatment together with increased total area of cells that were Kit^+^. By contrast, in salispheres cultured in simple media, Kit levels were low and did not increase significantly in the presence of Y27632. As the expansion in the epithelial area expressing Kit implies that Kit^+^ cells are proliferating, we performed ICC to detect Ki67 together with Kit. Ki67^+^/Kit^+^ cells expanded within the EPCAM^+^ area in salispheres grown in Sali media+Y27632 relative to Sali media alone, and much less so in simple media. These results imply that culture of salispheres in Sali+Y27632 increases the number of cells expressing Kit by stimulating proliferation of Kit^+^ cells and increasing levels of Kit in individual cells **(Fig 2d).** In Sali+Y27632, the percent of Ki67^+^/Kit^+^ cells is 59±5.7% **(Fig 2e)**, demonstrating that Y27632 can be used to increase the number of proliferative Kit^+^ cells in Sali media salisphere cultures.

**Figure 2:**
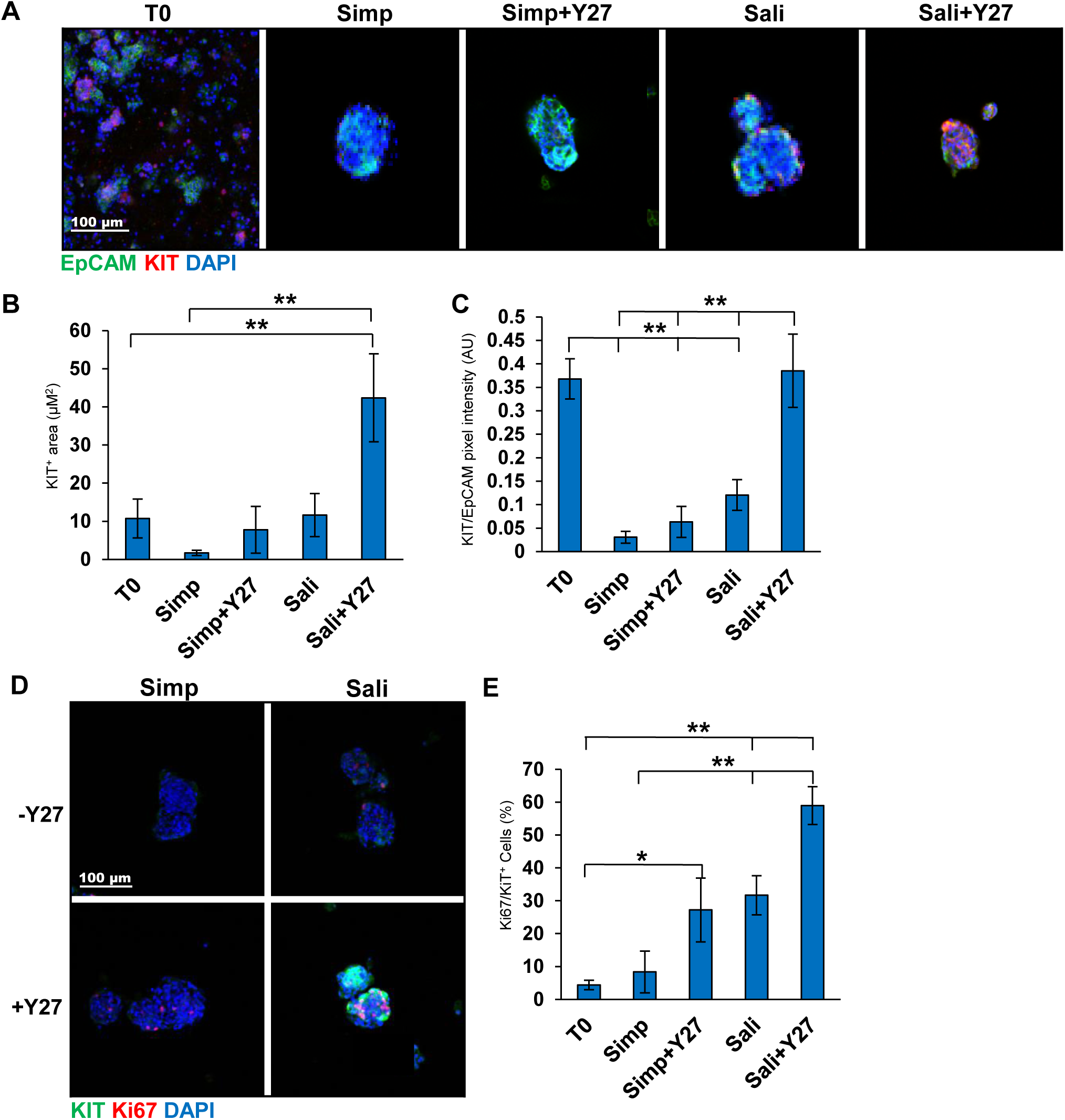
Y27632 treatment increases expression of Kit within salispheres grown in defined medium. A) ICC of Kit in salispheres grown in two medias +/- Y27632: EpCAM (green, epithelium), Kit (red, acinar/proacinar) with DAPI (blue, nuclei) in comparison to primary cells at time zero (T0). B) Quantification of Kit^+^ area in salispheres relative to EpCAM. C) Kit pixel intensity relative to DAPI. D) ICC of Kit (green) and Ki67 (red) in salispheres. E) Quantification of Ki67 in Kit^+^ cells (%). A-C N=7. D-E N=3. *p <0.05, ** p <0.01, One way ANOVA.

We also examined expression of the basal epithelial cell markers Keratin 5 (K5) in salisphere culture in the presence or absence of Y27632. In embryonic SMG K5 is expressed in the ducts, while in adult glands K5 is also expressed a subset of the myoepithelial cells. When K5 was examined by ICC in salispheres grown for 3 days with or without Y27632, K5 expression was robust in simple media with and without Y27632 **(Fig 3a)**. The K5+ area was significantly higher with Y27632 (7±3 vs 23±3) and significantly higher than T0 **(Fig 3b)**. Expression levels of K5 assessed by quantification of pixel intensity were also significantly higher with Y27632 (0.2±0.2 vs 0.3±0.02) and was also significantly higher than T0 **(Fig 3c).** Interestingly, salispheres grown in Sali and Sali+Y27632 showed little difference in K5 protein levels or salisphere area that is K5^+^. Keratin 14 (K14), which can dimerize with K5, also showed the highest levels in Ctrl+Y27632 media by quantification of pixel intensity (1.0±0.3 vs 0.6±0.1) **(Sup 1a).** In addition to the overall pixel intensity, the total salisphere area that was positive for K14 in the salispheres was also higher in simple media+Y27632 salispheres than in simple media salispheres. The K14^+^ pixel intensity of salispheres grown in Sali media was slightly greater than inSali+Y27632 (1.4±0.09 vs 0.9±0.06) **(Sup 1b-c)** similarly to what was observed with K5. Interestingly, K5/K14 proliferation in serum containing media was significantly greater than in the Sali media with or without Y27632, despite the fact that salispheres in Sali media were more proliferative **(Fig 3d-e, Sup 1d-e).** These results demonstrate that the simple serum-containing media supports and can expand the K5^+^ basal epithelial cell population.

**Figure 3:**
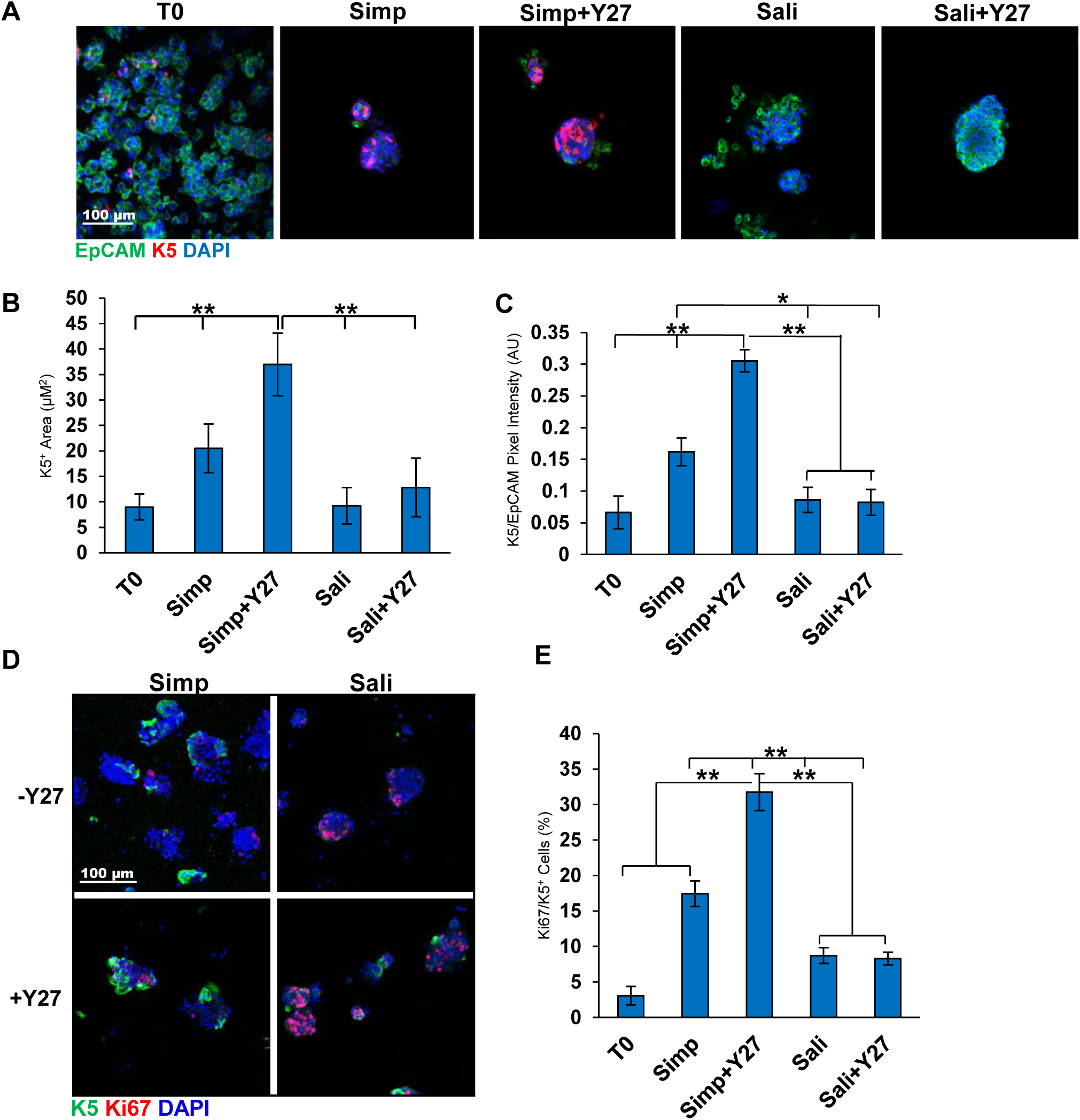
Y27632 treatment increases K5^+^ cell proliferation levels in salispheres grown in simple media. A) ICC of K5 in salispheres grown in two media conditions +/- Y27632: K5 (red), EpCAM (green), DAPI (blue). B) K5 pixel area relative to DAPI. C) K5^+^ pixel area relative to EpCAM. The area and intensity of K5 increases in salispheres grown in Ctrl media + Y27. D) ICC of K5 and Ki67 in salispheres. E) Quantification of Ki67/K5^+^ cells (%) reveals increased Ki67+ cells in Ctrl media + Y27. A-C: N=6, D-E: N=3 (Bl6 mice). *p <0.05, ** p <0.01, One Way ANOVA.

Since salispheres have also been reported to contain cells derived from Mist1^+^ secretory acinar cells (Varghese et al, 2019), we used Mist1^*CreERT2*^/ROSA26^*TdTomato*^ reporter mice to isolate salispheres and assayed for expression of reporter cells in both medias +/- Y27632 **(Fig 4a-b)**. Visualization of TdTomato revealed that Mist1^+^ cells contributed to salispheres formed in each of the four media conditions with no measurable change in TdT tomato in salispheres grown under Y27632 treatment any media condition, but increases in Sali media relative to simple media (**Fig 4c-d**). This indicates Mist1^+^-derived cells do not appear to be directly impacted by Y27632 but are enriched by Sali media relative to simple media.

**Figure 4:**
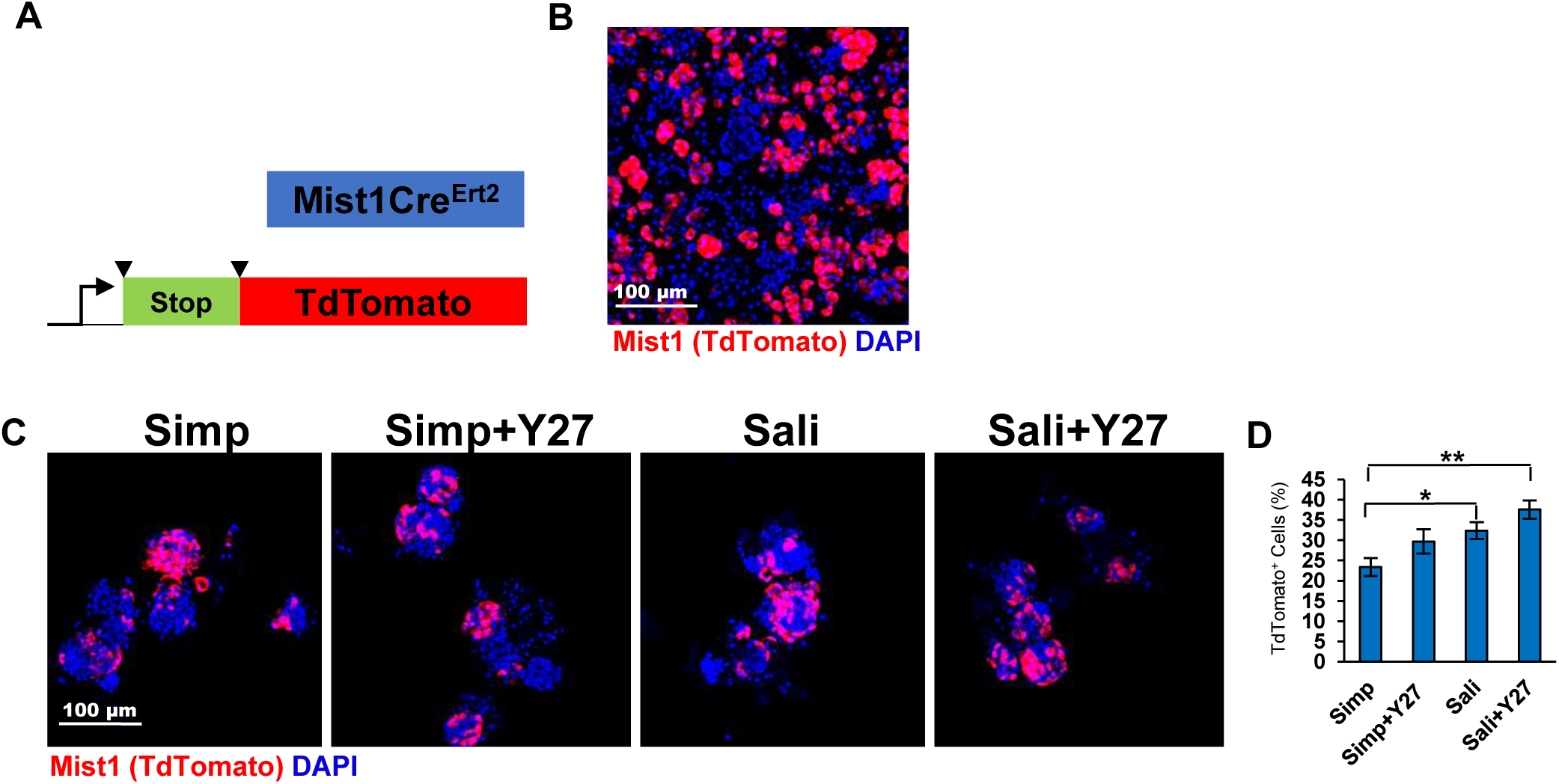
Mist1-derived cells are present in salispheres and are increased with culture in salisphere medium. A) Schematic of Mist1^CreERT2^; R26^TdT^ mice. B) ICC of Mist1^*CreErt2*^; R26^TdT^ mouse primary epithelial clusters at T=0. C) TdT expression in salispheres grown for 3 days. D) Quantification reveals that the average % of salispheres containing Mist1^+^ cells increases in Sali relative to simple media but is not significantly affected by Y27 treatment. N=3. *p <0.05, ** p < 0.01, One Way ANOVA.

### 3.3. Proacinar differentiation of salispheres in organoid culture is dependent on FGF2 and primary mesenchyme

Previously, we generated complex salivary organoids with robust, membrane localized AQP5 using E16 epithelial clusters cultured with their own primary mesenchyme cells (Hosseni et al, 2018 a,b). To determine if adult salispheres form AQP5^+^ budded organoids under organoid culture conditions and if salisphere culture conditions impact organoid formation, salispheres were produced using both medias ± Y27632, and for organoid formation salispheres were transferred to Matrigel ® and grown in simple media containing FGF2, E16 primary mesenchyme, or both (**Fig 5a**). Organoids derived from salispheres grown in simple media exhibited little AQP5 **(Fig 5d-e**). Organoids derived from salispheres grown in Sali media during the salisphere phase exhibited a modest but significant increase in levels of AQP5 when cultured with both FGF2 and E16 mesenchyme, that was further improved by Y27632 (**Fig 5b-c**). However, when compared to E16 epithelium derived organoids, organoids derived from salispheres grown in Sali media with Y27632 exhibited considerably less AQP5 intensity and AQP5/EpCAM^+^ area (**Fig 5f-h**). There was also a decrease in the total percentage of organoids (**Fig 5i**). These results indicate that proacinar differentiation in the salisphere-derived organoids is FGF-2 and mesenchyme dependent, but not as robust as proacinar differentiation in organoids derived from embryonic epithelium.

**Figure 5:**
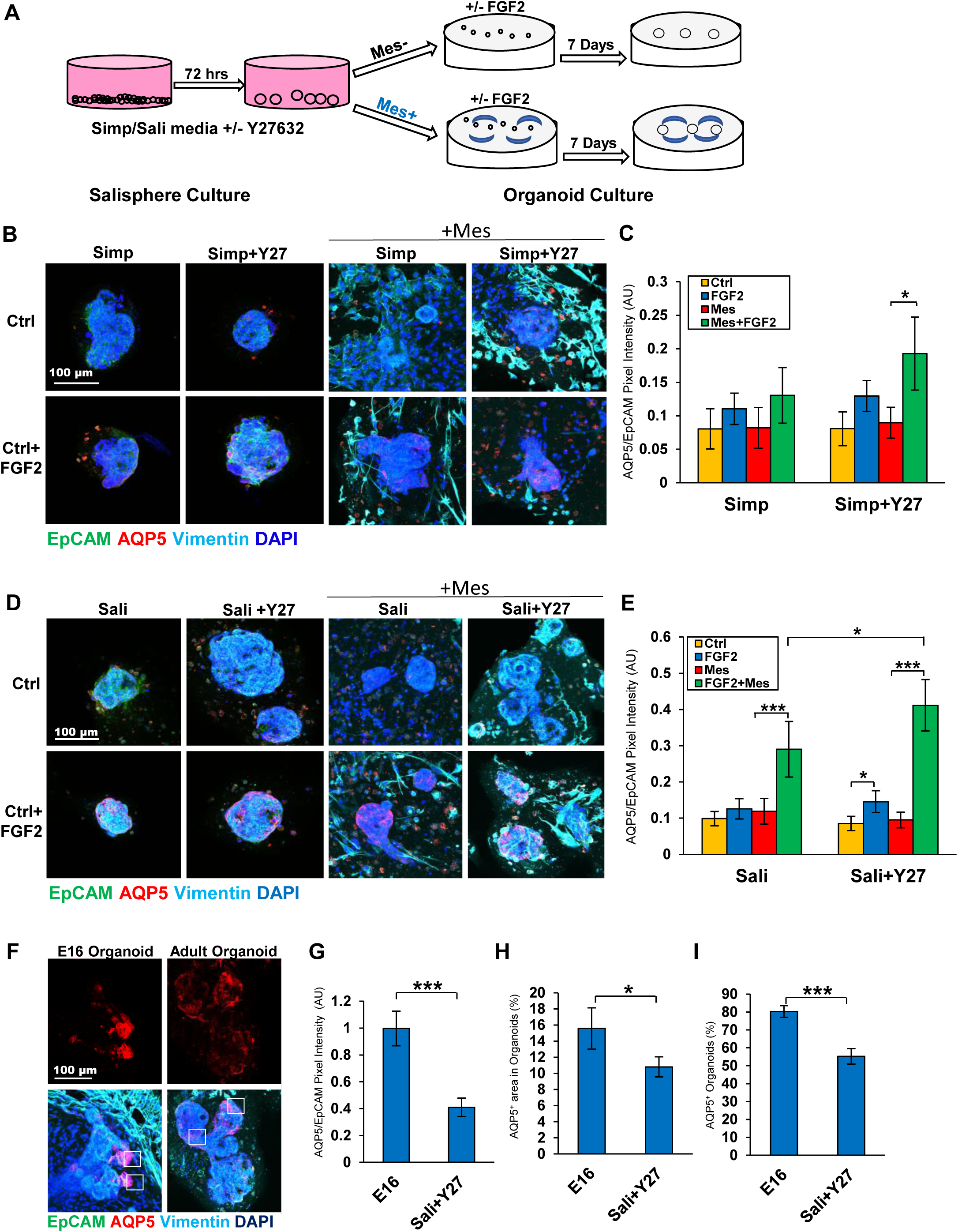
Salispheres grown with Y27632 generate proacinar salivary organoids. A) Experimental schematic of organoid experiments. B) ICC of salispheres grown +/- Y27632 in Simp media in the salisphere phase and in the presence or absence of primary E16 salivary mesenchyme cells (Mes) in Matrigel ® +/- FGF2 in the organoid phase. C) Quantification of salispheres of Ctrl vs FGF2 treated salispheres. N=5. *p <0.05, ** p <0.01, One Way Anova. D) ICC of salispheres grown +/-Y27632 in Sali media in the salisphere phase and in the presence or absence of primary E16 salivary mesenchyme cells (Mes) in Matrigel ® +/- FGF2 in the organoid phase. E) Quantification of AQP5/EpCAM intensity in organoids reveals that salispheres grown in Sali medium + Y27632 form proacinar organoids more effectively in the presence of Mes cells than salispheres cultured under other conditions. N=5. F) ICC of E16 derived organoids vs Adult Salisphere derived organoids grown in optimal conditions from B and C. White boxes indicate AQP5^+^ area. G) Quantification of AQP5 intensity of organoids relative to EpCAM. H) AQP5/EpCAM^+^ area. I) AQP5^+^ % organoids. Adult cells are less effective than embryonic cells at forming AQP5^+^ proacinar organoids. B-G: N=7. *p < 0.05, ** p < 0.01, ***p < 0.001, Students T-test (Sali+Mes+FGF2 vs Sali+Y27+Mes+FGF2), One Way ANOVA (C,E).

## 4. Discussion

Expansion of adult epithelial progenitor cells that can form secretory acinar cells is a goal for current applications in regenerative medicine. In this report, we demonstrate expansion of adult salisphere cultures to facilitate organoid formation in a second culturing phase. We demonstrate that treatment of adult SMG-derived salisphere cultures with the ROCK inhibitor, Y27632, can be used in combination with specific media formulations to expand specific epithelial cell populations. With salisphere medium in the presence of Y27632, we significantly expanded Kit^+^ cells by promoting cell proliferation. With growth in a simple serum-containing medium, inclusion of Y27632 in the media promoted expansion of K5^+^ cells. Organoids grown in culture that mimic specific aspects of whole organs can be used to examine cell potential. We used assays that we previously developed for embryonic SMG epithelial to examine the potential of adult salispheres to form complex salivary organoids with robust secretory proacinar differentiation. We demonstrate that the Y27632 can enhance the ability of adult salispheres to generate salivary organoids with secretory proacinar differentiation, although not as efficiently as organoids derived from E16 epithelium (Hosseini et al, 2018 a,b). As with the embryonic epithelium, we identified a requirement for FGF2 and the presence of mesenchyme cells for efficient organoid formation, suggesting that salispheres may retain the ability to further differentiate in vitro and in vivo in response to other signals.

A well-known function of Y27632 in cells cultured ex vivo is to promote proliferation as well as to ablate apoptosis in dissociated cells (Watanabe et al, 2007; Chapman et al, 2014; Sun et al, 2015; Croze et al, 2016; Zhang et al, 2011). ROCK inhibition by Y27632 has been also shown to delay senescence in the salivary gland epithelium (Lee et al, 2015). Although Y27632 has been included in salisphere culture in previous studies (Nanduri et al, 2014; Han et al, 2018), its direct role in salisphere formation was not previously documented. In this report, we demonstrate that Y27632 increases proliferation of cells grown in the salisphere medium. While we show that treatment of salispheres with Y27632 decreases apoptosis in serum-containing medium, there appears to be no difference in apoptosis reduction in salisphere medium, suggesting the presence of a survival factor in the salisphere medium that is lacking in the serum-containing medium. This suggests that in salisphere media, Y27632 has a more significant role in maintaining cell proliferation and in serum-containing medium has a more significant contribution to apoptosis, suggesting that crosstalk with other pathways stimulated by the media components modifies the effects of Y27632.

We here demonstrate that Y27632 can be used to differentially expand progenitor cell populations in specific media formulations. Treatment of salispheres with Y27632 increases proliferation of the K5/K14^+^ basal population in serum-containing medium and proliferation of the Kit^+^ cells in Sali medium. Y27632 has also been shown to increase progenitor cell markers in other model systems, including the limbal epithelium and in the neural crest (Miyashita et al, 2013; Kim et al, 2105). Mist1 is not unique to the salivary glands, as it is also expressed in the pancreas (Pin et al, 2001), but it is of interest as it is expressed in salivary epithelial cells that commit to the secretory acinar lineage. In development and in adult injury models, Mist1^+^ cells reproduce by self-duplication (Aure et al, 2015). Interestingly, the expansion of Mist1 was media-dependent and not Y27632-dependent. Nevertheless, our data is consistent with a prior report that salispheres are not exclusively ductally derived (Varghese et al, 2019).. However, the mechanisms driving proliferation of specific progenitor cell populations in specific media remains unknown and requires further investigation. Organoid assays have been used to grow a multitude of mini-organs in culture and are useful for exploring the contribution of specific cell types to forming organs. In the salivary gland, we previously reported that FGF2 in the presence of primary mesenchyme is able to stimulate a proacinar phenotype in organoids derived from E16 epithelium (Hosseini et al, 2018 a,b). We report here that the adult-derived salivary gland organoids are not identical to the embryonic organoids. Interestingly, while the organoids derived from salispheres grown in Sali media + Y27632 showed the highest levels of AQP5 amongst all other salisphere conditions, the overall levels of AQP5 expression was lower than that achieved by E16-derived organoids that we previously described (Hosseini et al, 2018 a,b). This suggests that the organoid capacity of the adult salispheres is limited relative to embryonic epithelium. Future investigation into niche requirements for adult salivary gland proacinar cells may lead to improved ex vivo growth conditions for adult cells for future application in regenerative strategies.

## Acknowledgments

The authors would like to thank Dr. Catherine Ovitt for generously providing us with the Mist1^CreErt2^ mice with permission of Dr. Stephen Konieczny. Finally, the authors thank Dr. Zeinab Hosseini for assistance with organoid culture methods.

## Funding

This work was funded by NIH grants: R56DE02246706, R01DE022467, RO1DE02246706 and by the University at Albany, SUNY.

AQP5: Aquaporin 5
K5: Cytokeratin 5
K14: Cytokeratin 14
SMG: Submandibular Gland
ROCK: Rho Associated Kinase
E16: Embryonic Day 16
T0: Time Zero
FGF2: Fetal Growth Factor 2
CC3: Cleaved Caspase 3
Salisphere media: Sali media
Simple Media: Simp media

## Notes

#### Summary of Updates

author list revised

