## Supplemental Figure 1 for "ROCK Inhibitor Increases Proacinar Cells in Adult Salivary Gland Organoids"

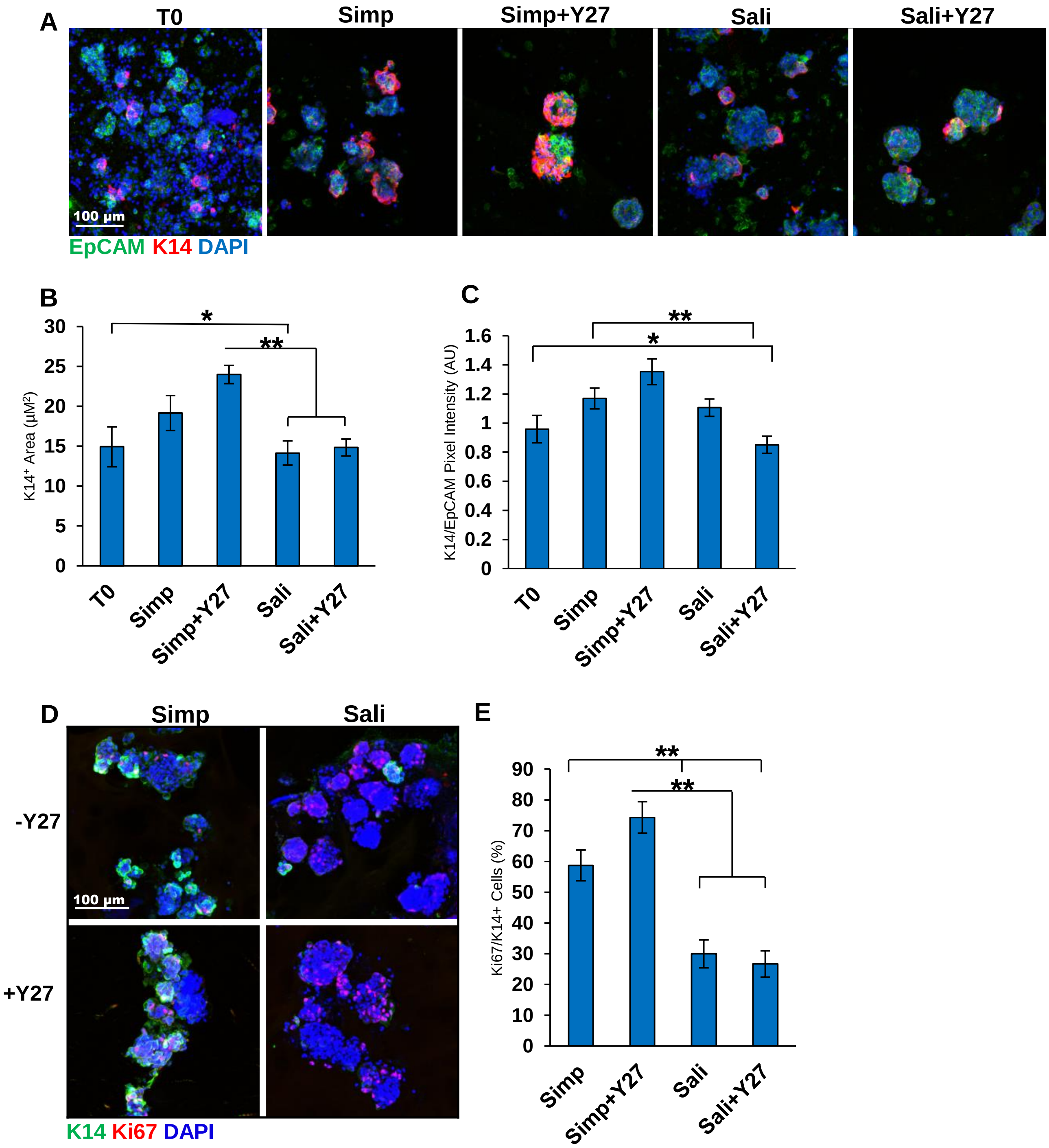


**Supplementary Figure 1: K14 increases in salisphere grown in serum-containing media.**

A) ICC of K14 in Salispheres. B) $K14^+$ pixel area relative to DAPI. C) $K14^+$ pixel area relative to EpCAM. D) ICC of salispheres with K14 and Ki67. E) K14/Ki67 proliferation. (%) A-C: N=6. D-E N=3 (Bl6 mice). *$p < 0.05$, ** $p < 0.01$, One Way ANOVA.
